# A Structural Antibody Benchmark of AlphaFold3 reveals Hallucinated Epitopes and a Bias for Orderness

**DOI:** 10.64898/2026.07.30.741792

**Authors:** Arnav Solanki, Neha Shree Maurya, Meaghan Ramlakhan, Rongbin Li, Wenbo Chen, Zhuhao Wu, Wenjin Jim Zheng

**Affiliations:** McWilliams School of Biomedical Informatics, University of Texas Health Science Center at Houston, Houston, Texas, United States; Appel Alzheimer’s Disease Research Institute, Feil Family Brain and Mind Research Institute, Weill Cornell Medicine, New York City, New York, United States

**Keywords:** AlphaFold3, Antibody, AI, Disorder, Epitope

## Abstract

AlphaFold3 has shown promise as a tool for predicting antibody-antigen binding, yet its performance across large datasets has not been fully characterized. In this study, 3401 experimentally validated antibody-antigen complexes were sourced from the Structural Antibody Database and screened alongside 23798 negative controls to benchmark AlphaFold3’s binding prediction capabilities. Confidence metrics including Predicted Aligned Error and Interface Predicted Template Modeling score were used to achieving a maximum recall of 53% at 100 inference seeds. Several factors were found to influence prediction accuracy: a notable bias was observed toward antibodies derived from X-ray crystallography structures versus those from electron microscopy, and positive prediction rates were found to decrease with increasing target protein size and surface area. In contrast, neither the amino acid composition or lengths of the complementarity determining regions, nor training data leakage were found to introduce significant bias. An innate false positive rate of approximately 3% was identified, with AF3 shown to hallucinate plausible binding interfaces across the surface of decoy targets while avoiding disordered regions. Epitope mapping using DockQ, epitope shift, and antibody displacement revealed that approximately 34% of false negatives retained the correct epitope location despite poor structural alignment, suggesting that conformation refinement tools could recover additional true binding predictions. These findings provide a comprehensive characterization of AlphaFold3’s strengths and limitations for antibody screening in computational drug discovery.

**Key Messages:**

- AlphaFold3 has a recall of 50% and an innate false positive prediction rate of 3%.
- False negative predictions can still feature the correct epitope despite poor RMSD.
- Factors such as disorder and target size impact accuracy.

## 1. Introduction

Antibodies are diverse proteins that the immune system has evolved to recognize pathogenic molecules. Antibodies work by uniquely binding target molecules and “tagging” them for the immune system to attack. The true binding target of an antibody is known as its *antigen*. Antibody-antigen binding is highly specific. Alongside the vast repertoire of antibodies in a host (at least a trillion unique antibodies per host! (5)), this ensures the immune system can identify virtually any possible antigen effectively. Their modular composition makes them excellent candidates for therapies for diseases such as cancer, and are also useful diagnostics tools for studying gene expression. However, generating and testing antibodies in a wet lab can be an expensive and laborious process.

This is where the recent generative-AI tool AlphaFold3 (AF3) has shown promising results (2). AF3 predicts protein structures and complexes from their sequences, and already boasts high accuracy in such modeling. These complex predictions often outperform traditional docking approaches, particularly when focusing on antibody-antigen interfaces. The overall structure of an antibody is mostly conserved and well annotated, and so AF3 has learned the basic blueprint for antibody-antigen binding. However, the immense diversity of antibodies and their high specificity poses a hurdle given the small structure data available for training in context of the search space (for example AlphaFold3 authors only analyzed 166 antibody-antigen interfaces (2)). A deeper understanding of how AlphaFold3 performs on a larger dataset, and one that also features negative controls is needed (13). That is the primary motivation behind our study: to study AlphaFold3’s performance when predicting antibody-antigen binding and decipher the various factors that influence it.

In this study, we benchmarked AlphaFold3’s performance on a larger dataset of antibodies derived from the Structural Antibody Database hosted by the University of Oxford. We observed AF3’s confidence is bimodally distributed when predicting positive versus negative binding, characterized by a recall of roughly 0.5. Metrics such as predicted aligned error (PAE) and interface predicted template modeling score (IPTM) showed high correlation with correctly predicting binding. We analyzed these results with structural information for a more refined validation. AF3 had a 3% false positive prediction rate where it hallucinated a wrong binding interface. 30% of the false negatives actually featured the correct binding site but displayed poor structural alignment. Factors such as the experimental method used to derive the structure (X-ray crystallography versus electron microscopy) and the disorderness and size of the target protein were observed to impact AF3’s predictions. In contrast, AF3 seemed unswayed by other factors such as the actual amino acids in the interface, or whether it had seen any particular antibodies before in training. This study demonstrates that AlphaFold3 is a powerful tool for antibody-antigen binding predictions with structural evidence, while highlighting the challenges that still remain in this domain of generative-AI models.

### 2. Background

Antibodies are symmetric Y-shaped proteins formed by the dimerization of 2 heavy chains and light chains (Figure 2). The heavy chains span the entire length of the antibody, forming the “trunk” of the Ab, whereas the light chains bind to the heavy chains at the “arms” of the protein. The top half of each arm of the antibody is known as the variable fragment F_V_ and is responsible for binding proteins. The F_V_ comprises 6 hypervariable loops that face the antigen when binding, collectively known as the Complementarity Determining Regions (CDRs). Each heavy chain and light chain contains 3 CDR loops. The remainder of the F_V_ is called the Framework Region. The large diversity of antibodies in a host is mostly a consequence of the variance of CDRs: they span various lengths and can consist of a variety of amino acids that lead to unique binding motifs for each antibody. On the other side of the interface, the true target of an antibody is called its antigen. The residues directly binding the antibody on the antigen are collectively called the epitope.

## 3. Methods

Here we present an overview of the methods we employed in this study; we have provided more technical details in Section A.

### 3.1. Data Mining

The Structural Antibody Database (SAbDab) tracks Protein Data Bank (PDB) files containing antibody-antigen complexes. We extracted the sequences and structures of 3401 non-redundant antibody-antigen complexes from SAbDab. We restricted the antibody sequences to the length of the variable fragment and annotated the CDRs using our custom definitions in Table 3. We attempted to fragment any long target proteins to isolate the subunit binding the antibody – only 133 long targets could be fragmented successfully. We recorded the full sequences of the remaining targets. Thus we yielded a positive control dataset with 3401 antibody-antigen complexes.

For each complex we tracked the type of experiment used for solving its PDB. We categorized the complexes into two batches: either M if the complex was captured in an electron microscopy (EM) PDB, or X if it was solved using X-ray crystallography. The M batch had 1342 complexes, and the remaining 2059 complexes formed the X batch.

Antibody databases generally do not report non-binding instances. As antibodies are specific, we reasoned that we could generate negative controls by screening antibodies against targets not reported as their true antigen. We implemented two types of negative controls:

1. One-to-One: We paired all positive control antibodies with random human proteins as non-binding targets – Each antibody to a unique target sampled from UniProtKB. We labeled these negative controls as the ℕ batch.
2. Many-to-One: We suspected that using only one unique negative target for each antibody was noisey and insufficient. To address this, we paired each of the 3401 antibodies against a fixed set of multiple negative targets. The proteins we selected were:

1. Batch A: *Rattus norvegicus* PVALB.
2. Batch B: *Drosophila melanogaster* CG6073.
3. Batch C: *Escherichia coli* araB.
4. Batch D: *Bos taurus* CLIC4.
5. Batch E: *Arabidopsis thaliana* DGR2.
6. Batch F: *Homo sapiens* CD274.

We tracked all PDBs for these targets to ensure none were also listed on SAbDab. 9 PDBs in our positive control contained CD274, so we removed those 9 antibody-target pairs from the F batch. The other 5 proteins had no PDBs containing antibodies.

Thus we generated a total of 23798 data points (3401 +(6 *×* 3401) − 9) as negative controls for our analysis. We used AF3 to predict the complex of all antibody-target combinations in our positive and negative controls over 10 seeds inference.

### 3.2. Epitope Mapping

Alongside the sequence data mentioned in Section 3.1, we also downloaded the PDB files of all antibody-antigen pairs. With these structures we were able to map the true epitope from the PDB and compare it with AF3’s prediction. Of course, we could only investigate the positive control predictions – the remaining data points were generated negatives and therefore had no meaningful epitopes. We used the following 3 metrics:

1. DockQ: We utilized this state-of-the-art quality measure to compare the predicted structures to the ground truth PDBs (4; 17). DockQ combines the root mean square difference (RMSD) of all the residues in the antibody-target interface and the fraction of contacting residues in both models to assess prediction quality.
2. Epitope Shift: This is the first of our two custom metrics for epitope measurement. It measures the locations of the predicted epitope with respect to the true epitope on the AF3 predicted target protein. A low epitope shift means that the residues in AF3’s predicted epitope are at the same location as the residues in the true epitope.
3. Antibody Displacement: This custom metric measures the change in the position of the antibody along the target surface in the prediction versus the ground truth. A low antibody displacement means that the predicted physical center of the antibody is close to the location of antibody in the original PDB.

We have provided more details on how we calculated these metrics in Section A.

## 4. Results

### 4.1. Overview of AlphaFold3’s performance

AlphaFold3 generates several scores such as Predicted Aligned Error (PAE), Interface Predicted Template Modelling (IPTM), and Ranking Score, that strongly indicate AF3’s confidence in a predicted complex. These metrics are generally correlated with each other (Figure 8). We picked PAE and IPTM as the crucial metrics for assessing binding prediction since they measure confidence specifically at the predicted interface rather than the whole complex.

In Figure 1 (B), the distribution of antibody-target complexes is bimodal – the two peaks in both PAE and IPTM suggest that AF3 treats most complexes as non-binding until a coherent signal is discovered, improving confidence. A low PAE score means that AF3 is confident about at least 1 pair of residues across the antibody-target interface. A high IPTM value means that AF3 is confident about the positions of residues in the antibody-target interface. These values do not guarantee high binding affinity – they merely convey that AF3 predicts the given antibody and target to bind as binary prediction. That is, they measure how confident AF3 is that the antibody and target bind, and not necessarily how strong they bind.

**Figure 1.**
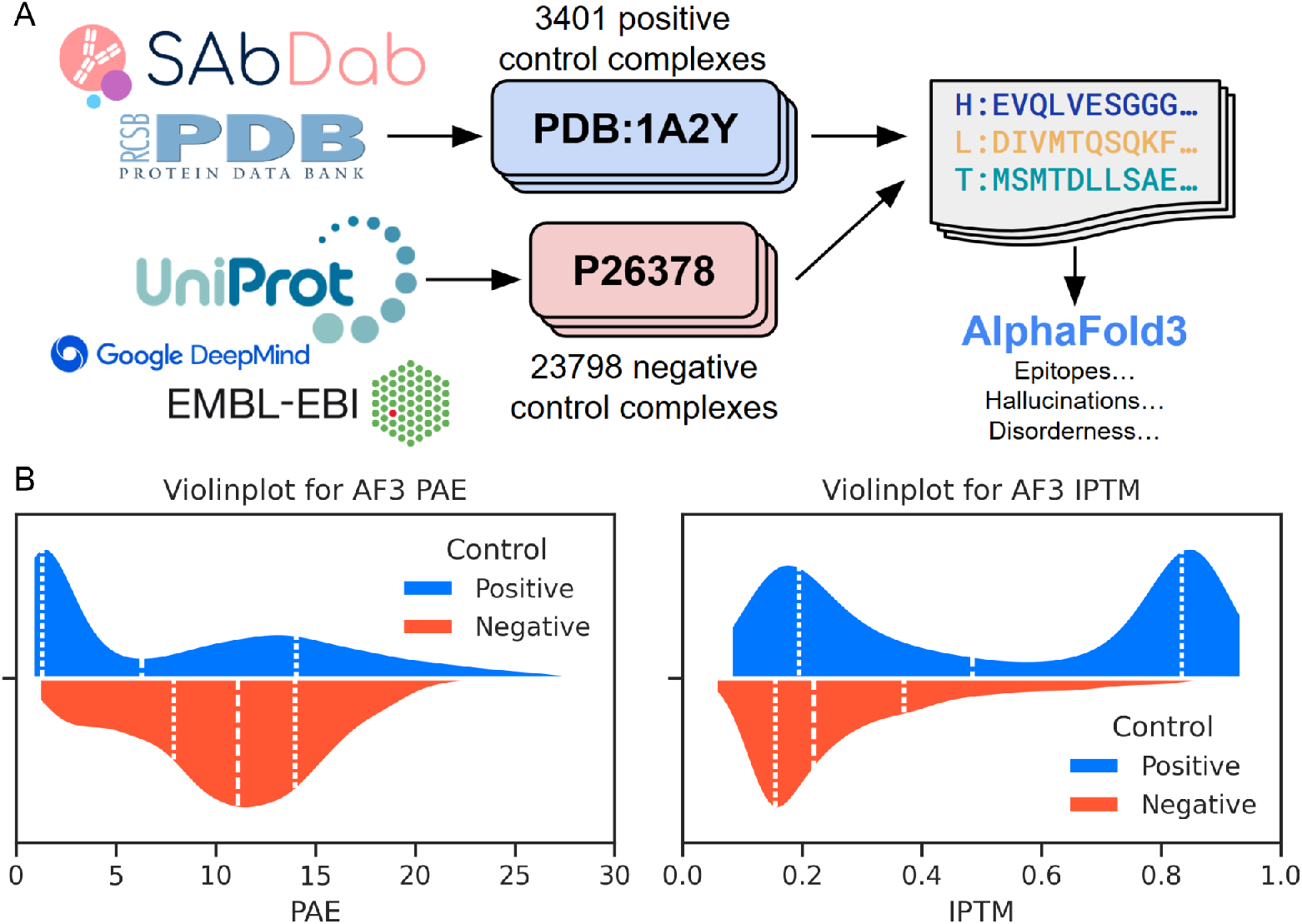
(A) An overview of our study. (B) Violinplots for the AlphaFold3 metrics: predicted aligned error (left) and interface predicted template modeling (right).

**Figure 2.**
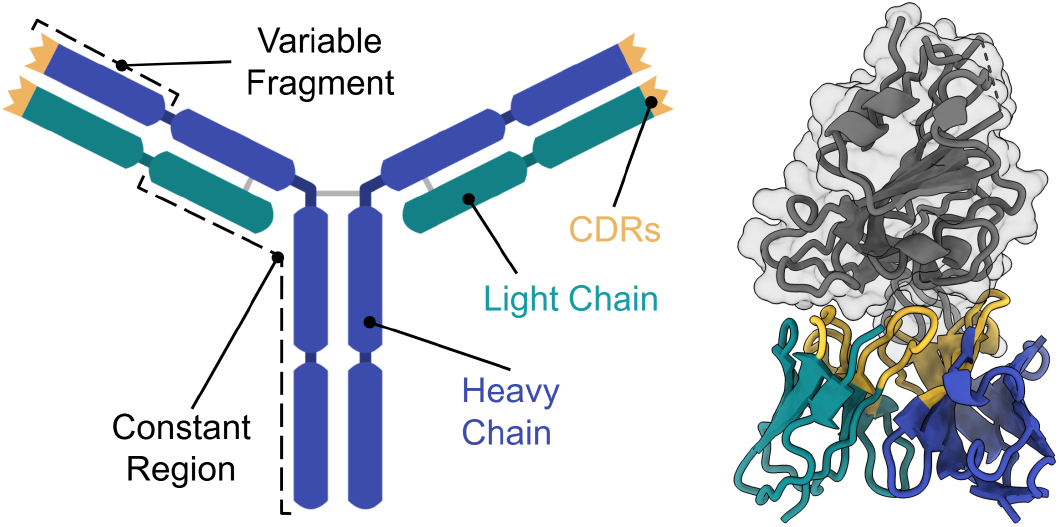
(Left) The general structure of an Antibody, showing its Y-shape and the variable and constant regions. (Right) The FV of an antibody as it binds its antigen (PDB: 8J1T). The heavy chains are colored blue, the light chains green, and the CDRs are yellow in both images. The antigen is colored gray.

Based on the distribution of the two metrics in Figure 1 (B), we classified a positive prediction based on the following criteria: A positive complex either had to have PAE less than 2.5 or IPTM greater than 0.7. The confusion matrix based on this criteria is laid out in Table 1. We yielded a total accuracy of 0.8997, recall of 0.4576, precision of 0.6367, and F1 score of 0.5325. Note that the high accuracy here is misleading; it is inflated by the large number of true negatives predicted from our negative control.

**Table 1.**
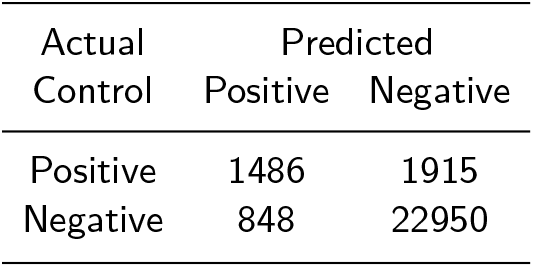
Confusion matrix of antibody-target predictions using AlphaFold3.

| Actual<br>Control | Predicted |  |
| --- | --- | --- |
|  | Positive | Negative |
| Positive | 1486 | 1915 |
| Negative | 848 | 22950 |

The PAE and IPTM scores distributed over all batches of antibody-target complexes (in Figures 9 and 10) show how positive and negative predictions in Table 2 are clearly distinct. Notably, the X batch of positive controls has a higher recall (0.52) than the M batch (0.30). AF3 has an evident bias towards antibodies derived from X-ray PDBs compared to EM PDBs. AF3 also predicts false positives across all negative control batches, suggesting it has an innate positive rate. Given these observations, we will delve deeper into the specific mechanisms that govern AF3 performance below.

**Table 2.** Antibody-Target Predictions using AlphaFold3 across various batches.

| Batch | M | X | N | A | B | C | D | E | F |
| --- | --- | --- | --- | --- | --- | --- | --- | --- | --- |
| Control | + | + | - | - | - | - | - | - | - |
| Number of Positives | 414 | 1072 | 176 | 115 | 86 | 66 | 110 | 193 | 102 |
| Number of Negatives | 928 | 987 | 3225 | 3286 | 3315 | 3335 | 3291 | 3208 | 3290 |
| Total Number | 1342 | 2059 | 3401 | 3401 | 3401 | 3401 | 3401 | 3401 | 3392 |

**Table 3.** Definition of each CDR using Chothia numbering.

| CDR | Chain | Chothia Residues |
| --- | --- | --- |
| CDR1 | Heavy | 26-35 |
| CDR2 | Heavy | 47-58 |
| CDR3 | Heavy | 93-102 |
| CDR4 | Light | 24-36 |
| CDR5 | Light | 46-56 |
| CDR6 | Light | 89-97 |

### 4.2. Factors that impact AlphaFold3 prediction

We investigated several factors that could have contributed to AlphaFold3’s middling performance. Here we highlight the factors that we discovered do impact antibody-antigen prediction.

#### 4.2.1. Size of the target protein

A simple observation from Tables 2 and 4 is that the length of the Many-to-One target protein impacts the positive prediction rate of AF3. The number of false positives in a Many-To-One batch (A through F) is inversely correlated to the length of that target protein (Pearson’s correlation = -0.682). This quick back-of-the-envelope calculation suggests that the larger a target protein, the lower the number of positives predicted to bind to it.

**Table 4.** PLDDT Distributions: Fraction of the residues of each negative control targets within the given PLDDT range Source: AlphaFold Protein Structure Database.

| Target used in negative batch | Length of protein | UniProt ID | PLDDT range |  |  |  | Average PLDDT | Fraction Disordered |
| --- | --- | --- | --- | --- | --- | --- | --- | --- |
|  |  |  | 0-50 | 50-70 | 70-90 | 90-100 |  |  |
| A (PVALB) | 110 | P02625 | 0.0% | 0.0% | 2.7% | 97.3% | 95.69 | 0.00 |
| B (CG6073) | 369 | Q9VBG6 | 0.0% | 0.3% | 6.0% | 93.7% | 95.69 | 0.01 |
| C (araB) | 566 | P08204 | 2.1% | 0.4% | 5.5% | 92.0% | 94.94 | 0.03 |
| D (CLIC4) | 253 | Q9XSA7 | 4.0% | 2.0% | 14.2% | 79.8% | 91.94 | 0.05 |
| E (DGR2) | 186 | Q94F20-2 | 0.0% | 4.3% | 14.0% | 81.7% | 93.19 | 0.02 |
| F (CD274) | 290 | Q9NZQ7 | 1.7% | 13.8% | 13.4% | 71.1% | 88.25 | 0.27 |

Source: AlphaFold Protein Structure Database.

Motivated by this observation, we investigated the impact of the length of the target protein on AF3’s binding prediction. We were particularly interested in AF3’s positive prediction rate and not necessarily on the true positive rate; thus we used both positive and negative controls. We binned all target proteins by their length into specific intervals as shown in Figure 3. The intervals were chosen to generate 10 equal-sized bins. We compared the distribution of PAE scores within each length bin. Clearly, the mean PAE score of negative predictions increases as the length of the target protein increases. Furthermore, the fraction of complexes predicted as positives decreases as the length increases, as shown in Table 5. The notable exception is bin 10, i.e. targets ranging from 500-2050. We believe this final bin is an outlier since the largest target in the ℕ batch was 500 residues long – all complexes in the final bin are positive controls only, hence the larger proportion of positives . Outside of this bin, it is clear that the length of target protein is inversely related to the positive discovery rate.

**Table 5.**
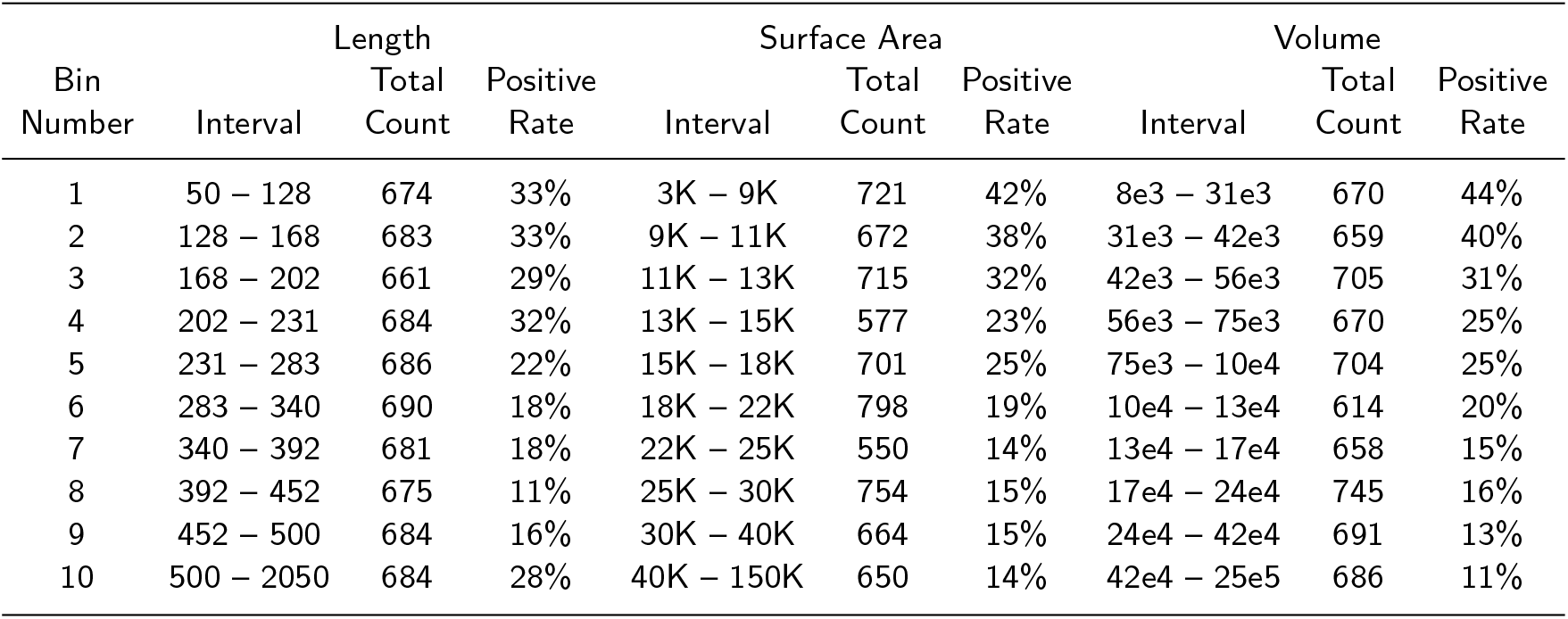
Target Predictions using AlphaFold3 across various batches.

**Figure 3.**
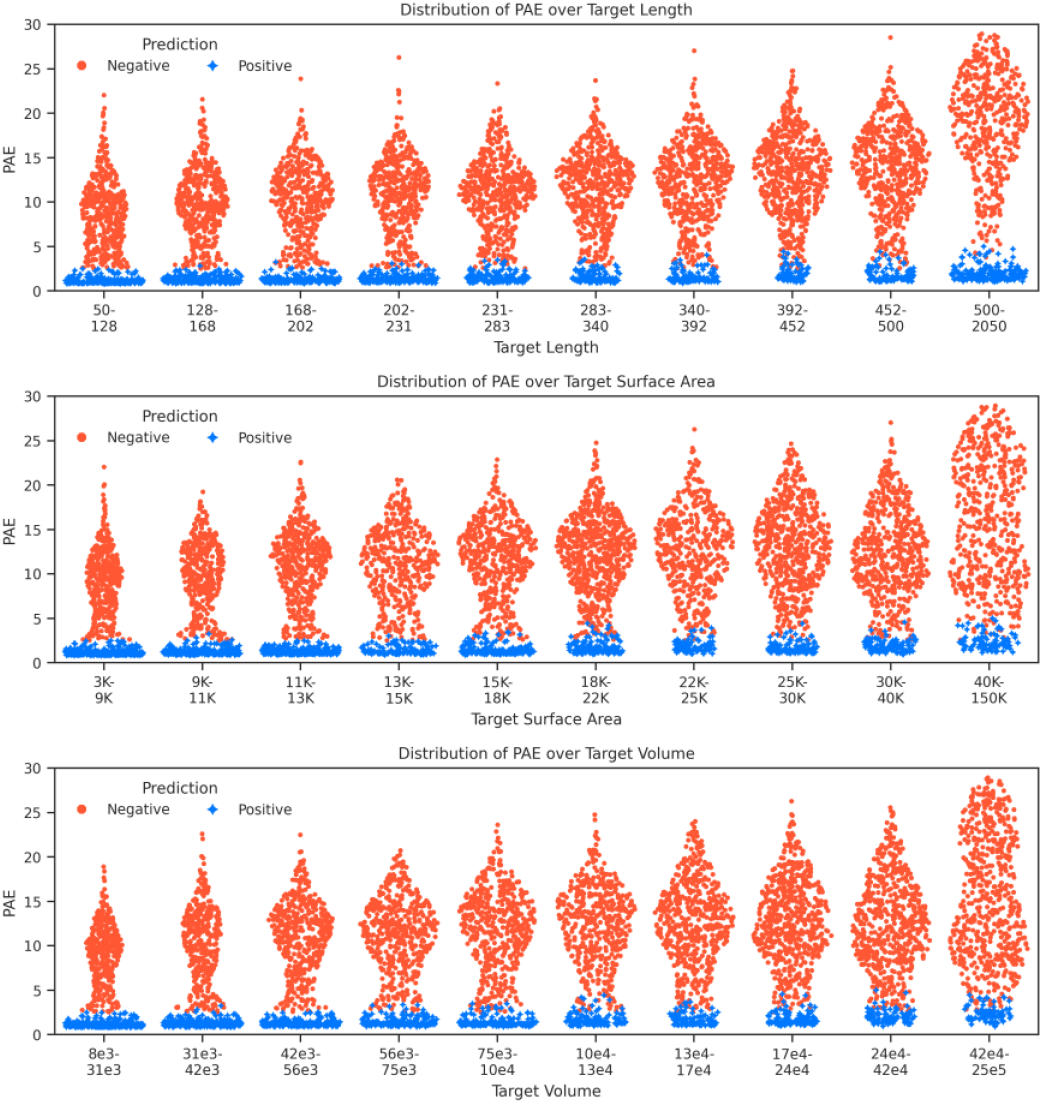
Sinaplot for the PAE score predicted by AlphaFold3 for an Antibody-Target complex versus Target measurements (Length, Surface Area, Volume). We binned the target parameters as shown on each x-axis. Complexes predicted as positives are shown in blue, while negative predictions are colored red. This analysis spans the M, X, and ℕ batches.

This trend of larger targets decreasing the positive discovery rate is even more prominent when analyzing the surface area and volume of the target protein in Figure 3. We used the Shrake-Rupley algorithm from BioPython to measure the surface area of every target protein, and the Convex Hull function from SciPy spatial algorithms to calculate the volume. The first 5 bins for the surface area and volume contain more positives (Table 5). Note that even though the ℕ target proteins were capped at a length of 500, their more disordered regions contribute to larger volumes and surface areas and hence the outlier of bin 10 in the length is not present. The main takeaway here is that the size and shape of target protein impacts how easily AlphaFold3 detects binding between the tested antibody and target. This is intuitive since larger proteins generate a larger search space for AlphaFold3 to predict the complex from.

#### 4.2.2. Disorderness of the target protein

The intrinsic order of an amino acid refers to its propensity to contribute to protein folding. Ordered regions of a protein are mostly populated by hydrophobic residues that form well-folded domains, often through structures such as alpha helices and beta sheets. Disordered regions tend to form peptide-like backbones that are flexible and dynamic in structure. It is well known that AlphaFold3 and similar protein folding tools have lower confidence when predicting disordered regions of a protein. For our analysis, we investigated the impact of the disorderness of a target protein on antibody binding.

We predicted the individual structures of all One-to-One targets, i.e. targets in the M, X, and ℕ batches. Here we left out the A through F batch since these were targets we had specifically hand-picked due to their ordered structures. Separately predicting the lone target structures allowed us to assess AF3’s confidence through the Predicted Template Modeling (PTM) score. As the true antigens in the positive control were derived from PDB structures, these proteins had higher PTM scores on average compared to the ones in the ℕ batch (Figure 4 (C)). However, positives were detected by AF3 across all values of target PTM, suggesting that AF3 is capable of identifying positives even for proteins with high disorder. The Pearson’s correlation of the target PTM with PAE for the positive control was -0.095, while for the negative control was 0.246.

**Figure 4.**
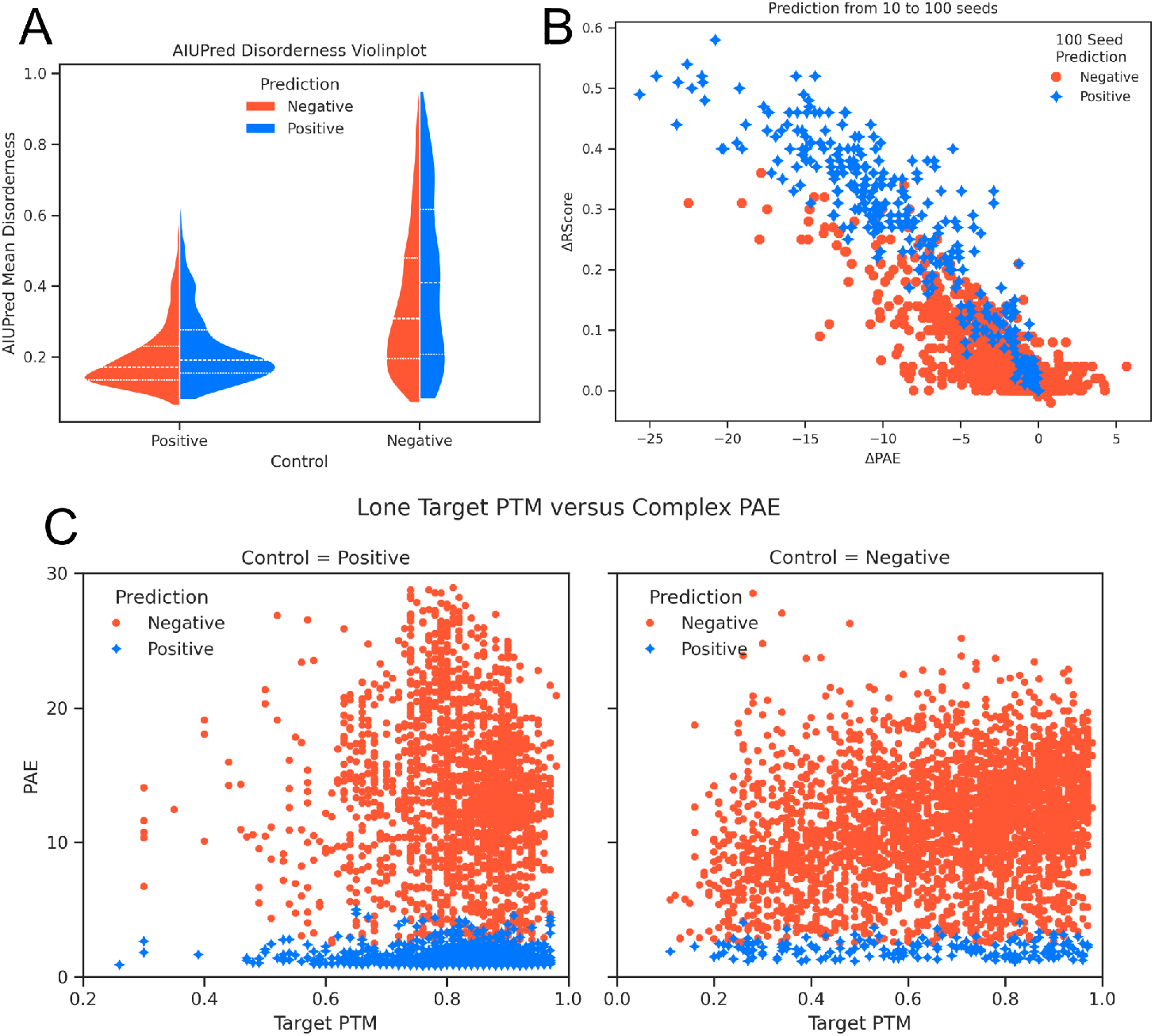
(A) Violinplot of target protein disorderness. (B) Scatterplot for improved AlphaFold3 metrics after 100 seed inference on the 1915 false negatives from 1. The ΔPAE and ΔRScore are measured as the PAE and Ranking Score from the 100 seed inference minus the corresponding value from the 10 seed run (respectively). Blue markers represent complexes that cross our criteria in Section 4.1 and become true positives after the 100 seed inference; the complexes that still remain as false negatives are colored red. (C) Scatterplots of PTM of the Target alone versus PAE of the complex. Left contains the M and X batches, while the right contains the negative control ℕ. Left corr = -0.095, right corr = 0.246

We also used the state-of-the-art AIUpred, a sequence-based machine learning tool, to predict the disorderness of every residue in our targets (12; 11). The mean AIUpred disorderness of a lone target was highly correlated with the disordered fraction reported by AF3 for that target (Pearson’s correlation = 0.8 across the M, X and ℕ batches). AF3 detects positives (both false and true as in Figure 4 (A)) on target proteins of all fractions of disorderness. We used a Mann-Whitney U test to confirm that the distributions of mean disorder across the positive and negative predictions was statistically consistent. For the negative control, the two prediction classes were not statistically distinct (p-value = 0.001 but with a mean difference of 0.07). For the positive control we calculated a stronger p-value of 5.6 *×* 10^−16^, but the mean difference between the positive and negative predictions was a mere 0.02. These values suggest that though there may be a minuscule difference in the disorderness of what AlphaFold3 classifies as a positive or negative, it does not have a large enough impact to show up when comparing it over the broad spectrum of possible proteins and how disordered they are. Furthermore, the false positives in more disordered negative targets (the tail on the right violin in Figure 4 (A)) are actually a consequence of the orderness of the positive control (no long tail on the left violin in Figure 4). Antigens extracted from PDBs are too ordered!

There is one important caveat however: Disordered regions are stable, functional components of several proteins, and can form strong epitopes on their own. Since their dynamic state can be difficult to capture in crystal structures, AF3 has low confidence when making predictions involving disordered regions. All of the antibodies in our analysis were extracted from the PDB, and therefore likely targeted ordered epitopes in their true antigen (we have not confirmed that every epitope in the positive control is fully ordered). The sample of antibodies we tested in our analysis is not representative of the whole set of naturally evolved antibodies.

Despite the observation that disorder of the target protein did not drastically influence positive prediction rate, protein disorder is a crucial factor that AlphaFold3 is not fully robust at predicting around. As noted in Section A.1.3, CD274, the target protein for our F batch of negative control, has a prominent disordered N-terminus. AlphaFold3 reports that 27% of the protein is disordered, and the last 50-60 amino acids of the protein are all predicted as disordered residues by AIUPred. This region can be spotted (in Figure 5) as the peptide sticking out from the rest of the folded protein. When we superimposed all antibodies that were falsely predicted to bind to it, we observed that none of the antibodies targeted the disordered region. This suggests that when AlphaFold3 tests an antibody against a potential target, it ignores the disordered regions of that target and only focuses on discovering epitopes on the folded regions.

**Figure 5.**
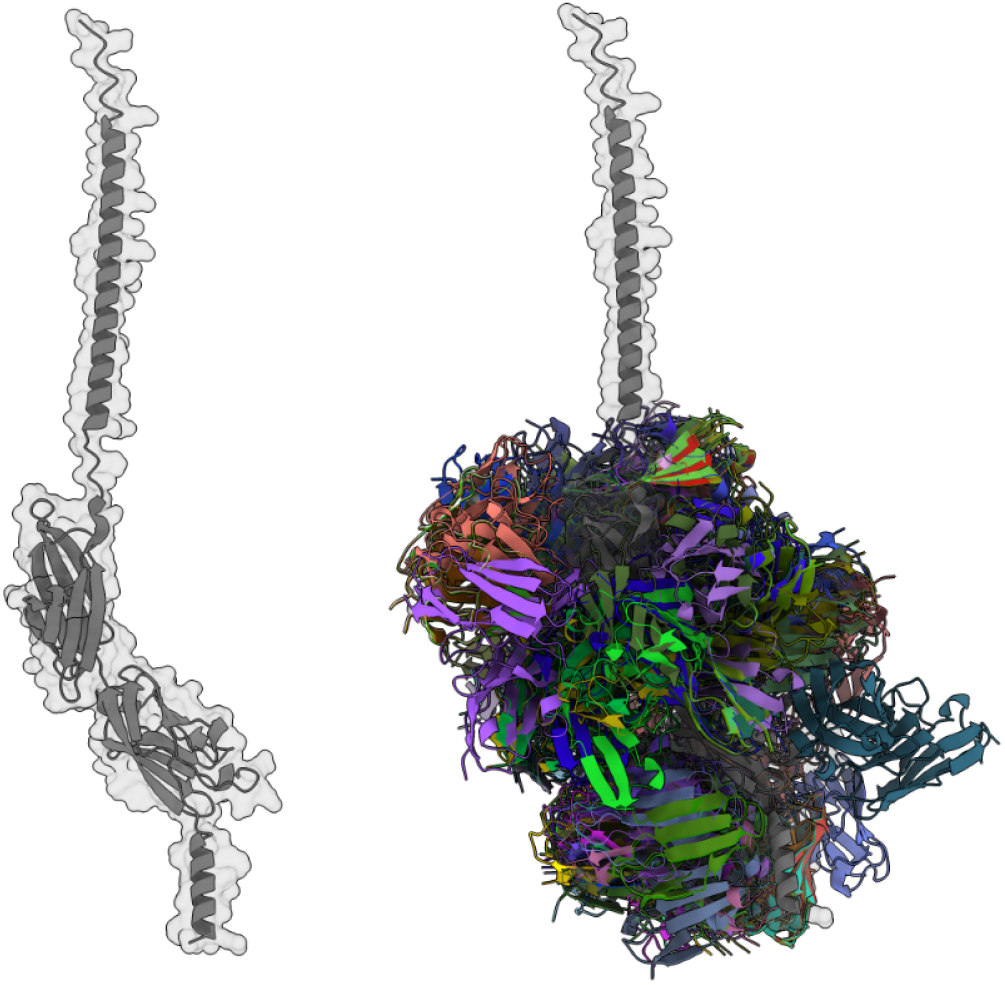
(Left) CD274, the target protein used in the negative control batch F. The long alpha-helix jutting out on the top forms the disordered N-terminus. (Right) All false positive antibodies targeting CD274, superimposed together. None of these antibodies were predicted to bind to the disordered region.

#### 4.2.3. Seeds for Inference

As noted by the authors of AlphaFold3, increasing the number of seeds during inference can increase the probability of discovering a positive antibody-target prediction (2; 15). We ran all 1915 false negatives (in Table 1) through AF3 again for a 100 seeds. AF3 reports the best model out of the results of multiple seeds by ranking score. Figure 4 (B) shows the improvement in AF3’s confidence. In most instances the ranking score of the complex increased. Correspondingly the PAE score improved too, and several complexes crossed our criteria for positive binding. Interestingly, we observed a gain in PAE in some instances (data points with positive ΔPAE in Figure 4 (B)) despite an improvement in ranking score. This means that sometimes AF3’s confidence improves with more seeds, and it then claims (with more certainty) that the given complex is not binding. A potential means of addressing this issue would be to manually choose the model with the best PAE score out of the results of multiple seeds instead of letting AF3 choose it.

A total of 344 antibody-target complexes that were previously not detected now crossed our binding criteria true positives after inference with more seeds. Accounting for this gain on the original 10 seed numbers, we got 1808 true positives and 1593 false negatives. This comes out to a recall of 53%. However, running AlphaFold3 for more seeds is computationally expensive. It is possible that running the remaining false negatives for 1000 seeds could improve the positive hit rate further, but this would be beyond the scope of our investigation.

### 4.3. Factors not influencing AlphaFold3 prediction

We observed several factors that did not seem to influence AlphaFold3’s predictions for antibody-antigen binding.

#### 4.3.1. Data Leakage

AlphaFold3 was trained on protein structures in the PDB. Since the antibodies in our positive control were derived from PDB files listed in SAbDab, there was a possibility of a bias towards antibodies AlphaFold3 had already “seen” in training in our analysis. To investigate this, we compiled the set of all PDBs used to train AF3 (a total of 195858 CIF files) and tracked all SAbDab PDBs that we extracted antibodies from. We intersected these two sets, and labeled all antibodies in our control that had been derived from an intersecting PDB as “leaked” antibodies. Out of the 3401 positive control antibodies, 2323 had leaked, i.e. AF3 had been trained on them, while 1078 were unseen to AF3 in training. Significantly more antibodies had leaked from PDBs captured using X-ray crystallography in comparison to PDBs solved using electron microscopy – this is not surprising since the former has been the more popular experimental method up until recently. Out of the 1342 M antibodies, 652 had leaked and 690 had not. For the 2059 X antibodies, 1671 had leaked while 388 had not. The differential prediction of AlphaFold3 on antibodies that had leaked is shown in Figure 6 (B). For each batch, the set of leaked versus not leaked antibodies could not be statistically distinguished using a Mann-Whitney U test on their PAE score (the lowest p-value observed was for the C batch at 0.016).

**Figure 6.**
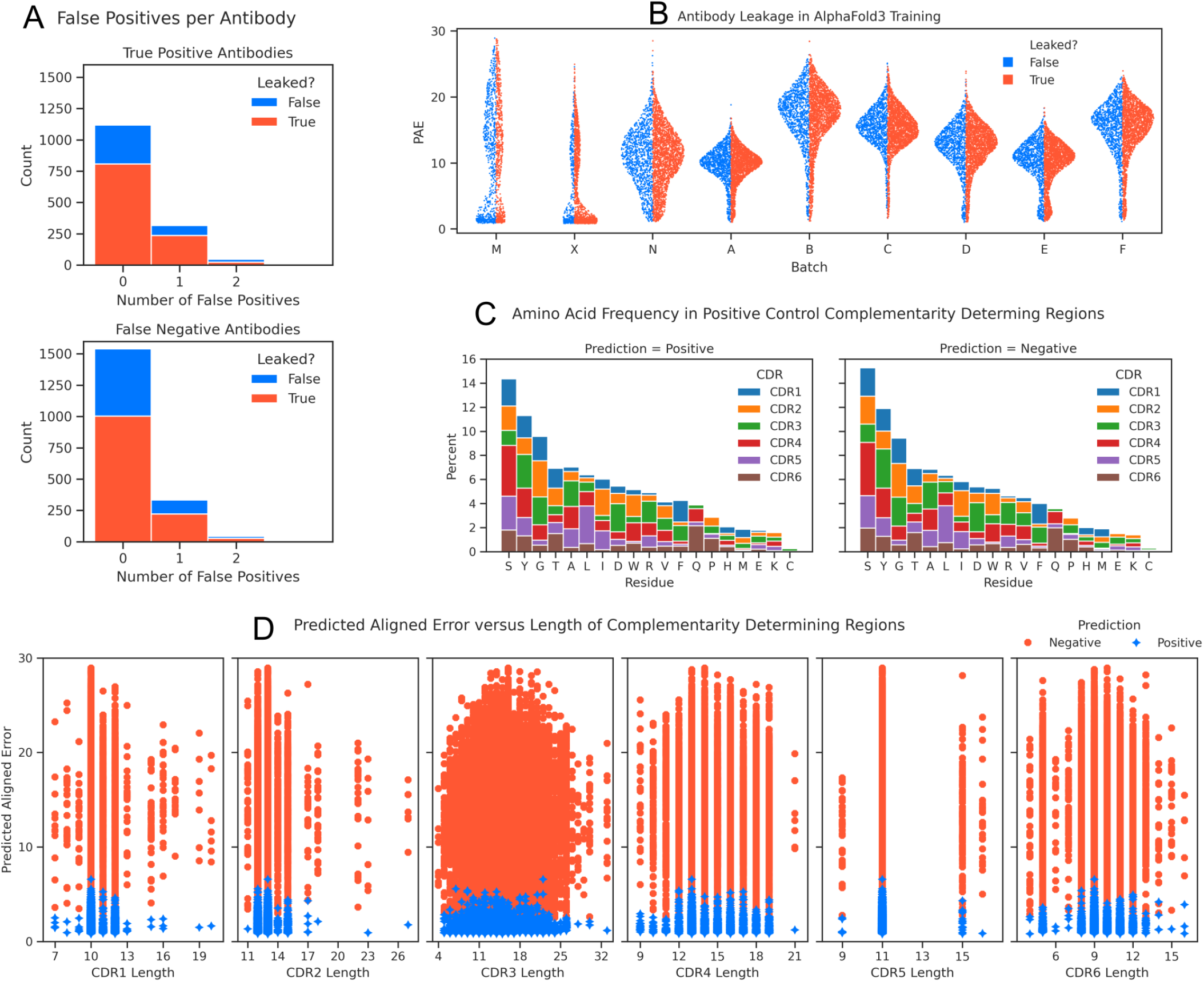
(A) Histogram of the number of false positive predicted in all batches. The top graph shows antibodies AF3 could predict the true antigen for while the bottom shows the false negatives. (B) Sinaplots for AlphaFold3 PAE while tracking leaked antibodies. The data is shown across all batches. Not leaked antibodies are shown in blue on the left, leaked antibodies in red on the right. (C) Frequency for each amino acid in all 6 CDRs of the positive control antibodies. The histograms are split by AlphaFold3 prediction. (D) AlphaFold3 PAE versus the length of each CDR for all antibodies. Positive predictions are colored blue and negative predictions are red (from all batches).

While AF3 does not allow users to “guide” antibody-target prediction by supplying information of a potential epitope in the input, it does provide the option of using a template PDB file to fold any individual chain. Note that these templates do not influence the binding prediction directly, and at best improve AF3 by giving it a good backbone to model either the antibody or target. AF3 is forced to compute the binding interface of every input from scratch. Since we observed leaked false negatives and true positives that had not leaked, it is likely that AF3 has no overwhelming bias towards antibodies it has already been trained on.

#### 4.3.2. Antibody Favoritism

Given the number of antibody-target complexes predicted as positive binders, we checked if AF3 was biased towards any particular antibody in predicting positives. Our concern here was that AF3 might treat certain antibodies as “promiscuous” and would latch them on to any available target surface screening. For each of the 3401 antibodies in our positive control, we tracked how many negative targets it falsely bound in our screening (shown in 6 (A)). A majority of the antibodies did not bind to any negative target or only bound to one negative target. Less than 10 antibodies falsely bound to 3 negative targets. Across antibodies that were predicted to bind to their true antigen and those that remained false negatives, the trend of false positive rates were similar (a Pearson correlation of 0.997). This means that a true positive prediction for a given antibody does not entail that that antibody is more prone to false positive predictions. When AF3 falsely predicts an antibody to bind to a random target, it truly identifies (or rather, hallucinates) a complementary epitope for that given antibody rather than just assuming that antibodies are “sticky” molecules that have to bind any available target.

#### 4.3.3. Complementarity Determining Regions

We investigated if any particular amino acid present in any CDR in our positive control antibodies biased AlphaFold3 towards predicting binding. Comparing the two histograms (split by AF3 prediction based on our criteria in 4.1) in Figure 6 (C), the distribution of all amino acids in all CDRs appear statistically equivalent for positive and negative predictions. Outside of Tyrosine (T) and Alanine (A), which have nearly equal frequencies in both classes, even the order of frequency of each residue is preserved. AF3 can accurately model the binding characteristics of CDRs with diverse sequences, and is not biased by the presence of any particular amino acid when predicting positives.

The length of the CDRs themselves could also be a factor in modeling binding. A longer CDR is more flexible; this could allow AF3 to “overfit” it on the surface of a target protein during screening. A long CDR can also be unstable due to higher disorderness, and this could lead to AF3 losing confidence when predicting on it. We measured the lengths of all CDRs in our dataset, and compared it with the PAE score of that antibody when tested against its true antigen as shown in Figure 6 (D). Note that each CDR spans different length intervals – CDR3 in particular is the most diverse in antibodies. In Figure 6 (D), a wide range of PAE scores were predicted for all lengths of CDRs. We did not observe any significant pattern in how a particular CDR of any certain length could bias AlphaFold3 (Pearson’s correlation for each CDR length with PAE was -0.01, 0.005, 0.108, 0.023, -0.001, 0.022). This suggests that when AF3 has high confidence in a given antibody-target complex’s binding, it is not caused by any spurious artifacts of any CDR. Rather, high confidence indicates that AlphaFold3 identified the epitope and antigen-binding site of antibody as truly complementary to each other.

### 4.4. Epitope Mapping

DockQ is a popular metric for scoring predicting protein interfaces, and it unsurprisingly correlates with the PAE of positive control antibody-complexes (Pearson’s correlation = -0.771). A high DockQ score indicates that the predicted model’s interface has minimal RMSD with the ground truth structure. This is evident in Figure 7 (A), where DockQ cleanly differentiates true positives from false negatives. The epitope shift score measures the change in the centroid of the epitope in the ground truth to the predicted model. It has a Pearson’s correlation of 0.641 with AlphaFold3 PAE over all positive control complexes. While this is lower than DockQ’s correlation with PAE, it is due to epitope shift measuring the location of the epitope and not necessarily the quality of the binding interface. It has a Pearson’s correlation of -0.592 with DockQ (Figure 7 (B)). From Figure 7 (A), it is clear that a positive prediction requires a low epitope shift. The antibody displacement measures the change in the position of the antibody in the prediction relative to the original location on the target protein. It has a Pearson’s correlation of 0.696 with PAE, and -0.743 with DockQ (Figure 7 (B)). Like the epitope shift score, a small antibody displacement suggests that the antibody was predicted to bind around the true epitope.

**Figure 7.**
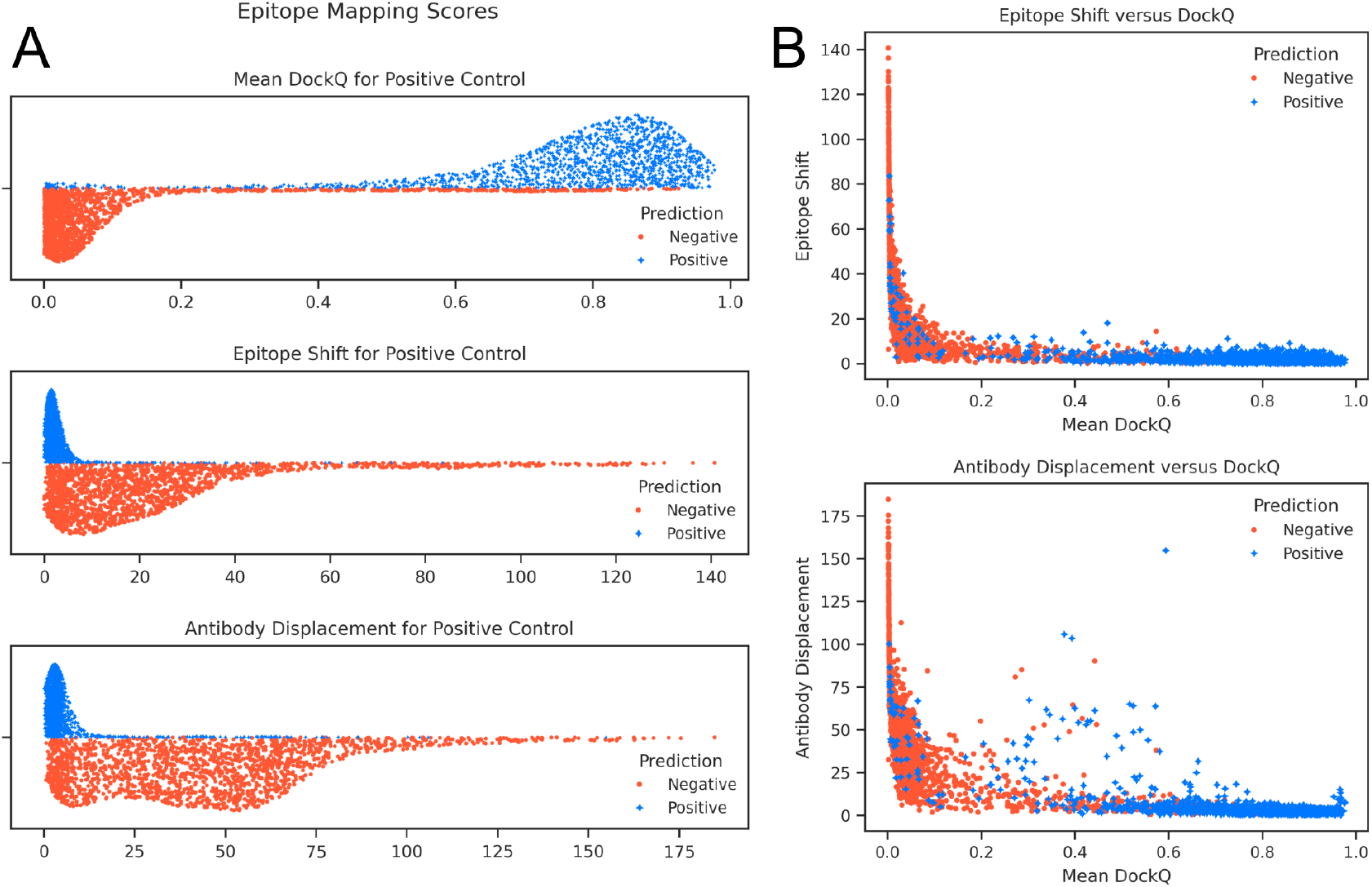
(A) Sinaplots for all 3 Epitope measurement scores (DockQ, Epitope Shift, and Antibody Displacement). High DockQ scores indicate a true positive complex prediction with a high quality interface. Low shift and displacement scores identify instances where the correct epitope was discovered in the predicted complex. (B) Scatterplots of Epitope Shift versus DockQ (top) and Antibody Displacement vs DockQ (bottom) over the positive controls.

**Figure 8.**
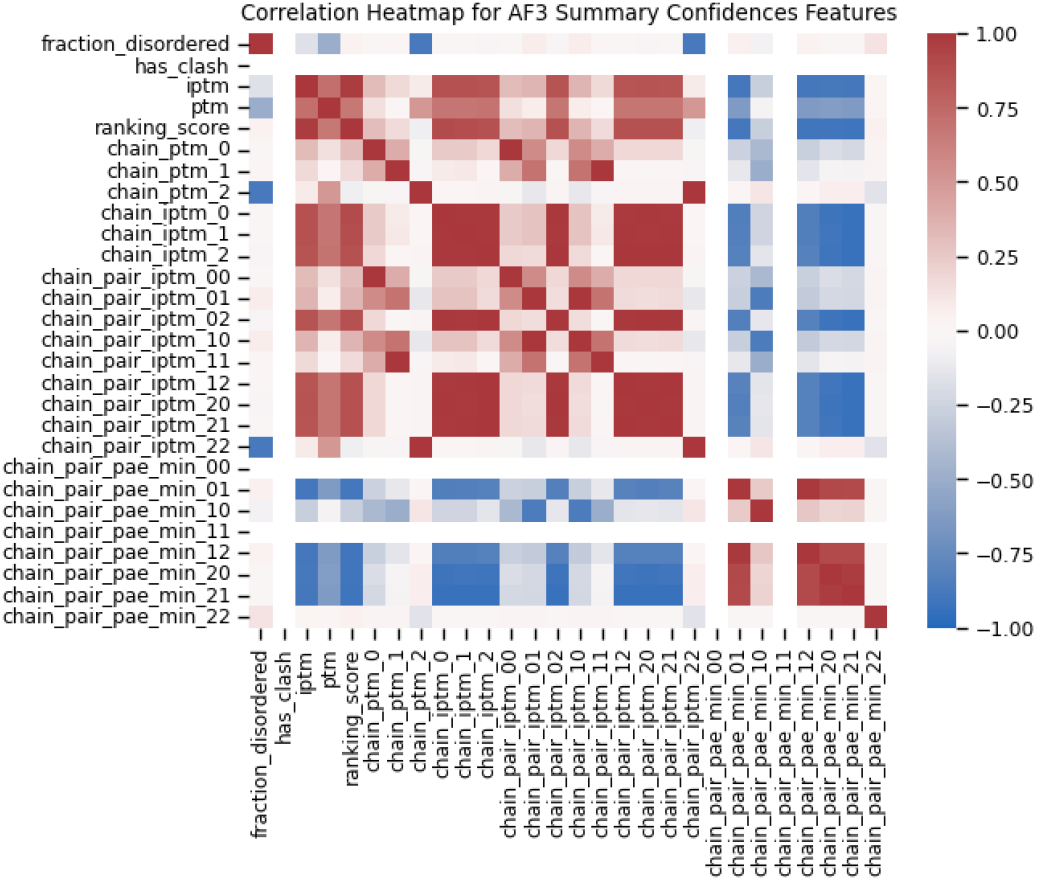
A correlation heatmap of the AlphaFold3 summary confidence scores across all 27199 antibody-target predictions.

**Figure 9.**
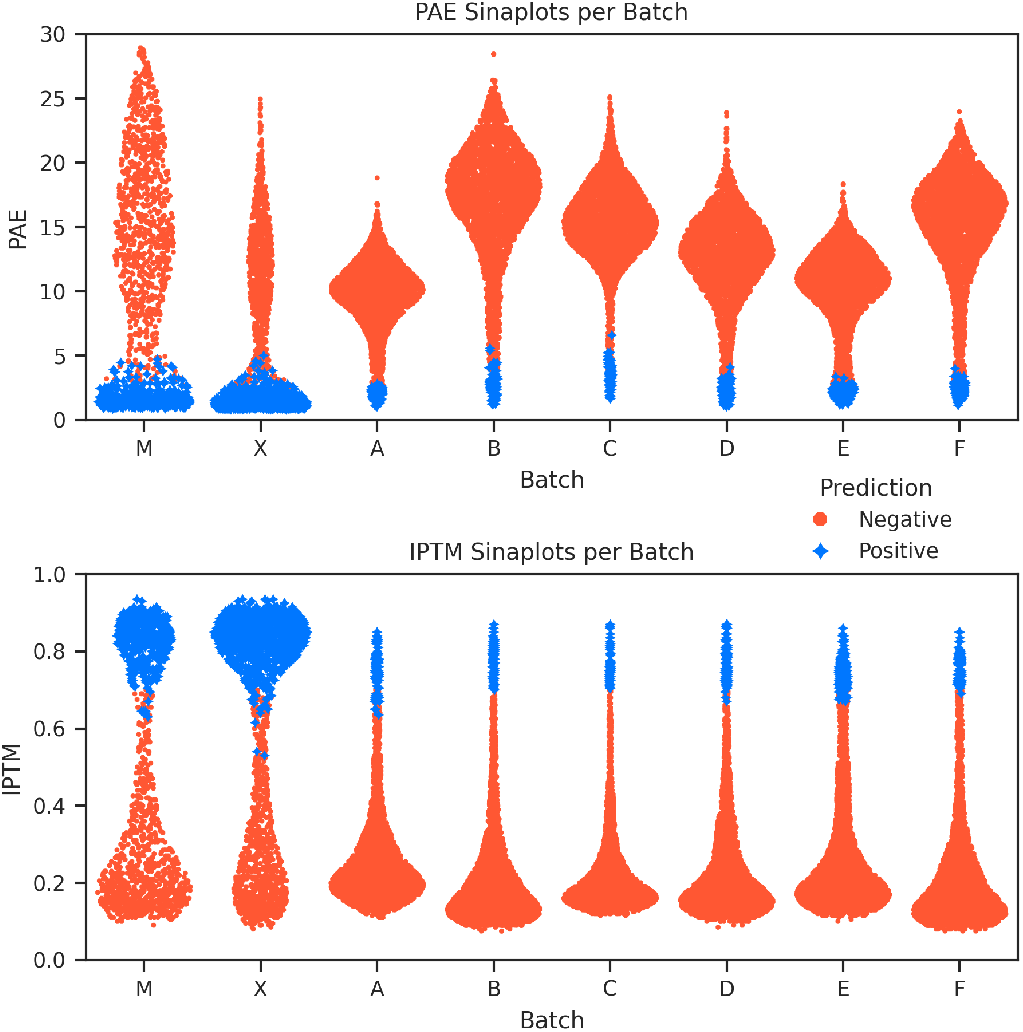
Sinaplot for PAE and IPTM across the different batches tested: M and X are positive controls, ℕ, A, B, C, D, E, F are negative controls

**Figure 10.**
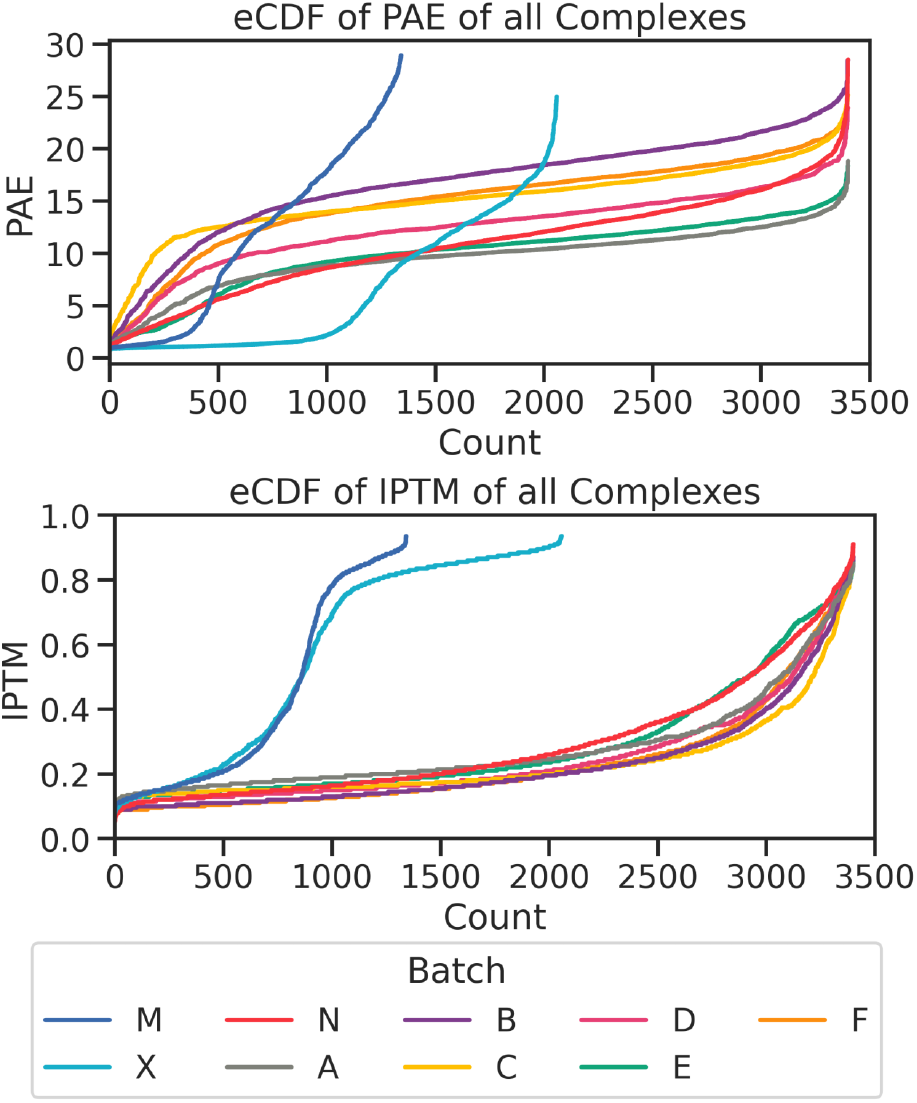
Empirical distribution functions (eCDFs) for AlphaFold3 PAE and IPTM across all batches. The x-axis represents the number of complexes. It is clear that the several positive controls in M and X have higher confidence through the low PAE and high IPTM scores. Clearly more X complexes were predicted as positives.

141 of the predicted true positives (from Table 1) had a DockQ score less than 0.5. However, a low DockQ score does not necessarily mean that the predicted antibody-target complex features the wrong epitope. A poorly folded protein or the wrong conformation of local residues at the epitope can increase the RMSD between the two structures even if the correct epitope was discovered. This is why we incorporated our additional epitope measuring algorithms in this analysis. Of these 141 complexes, 87 still had an epitope shift less than 10; we provide 3 examples out of these in Figure 11. Only 54 of the 1486 predicted true positives had largely incorrect epitopes.

**Figure 11.**
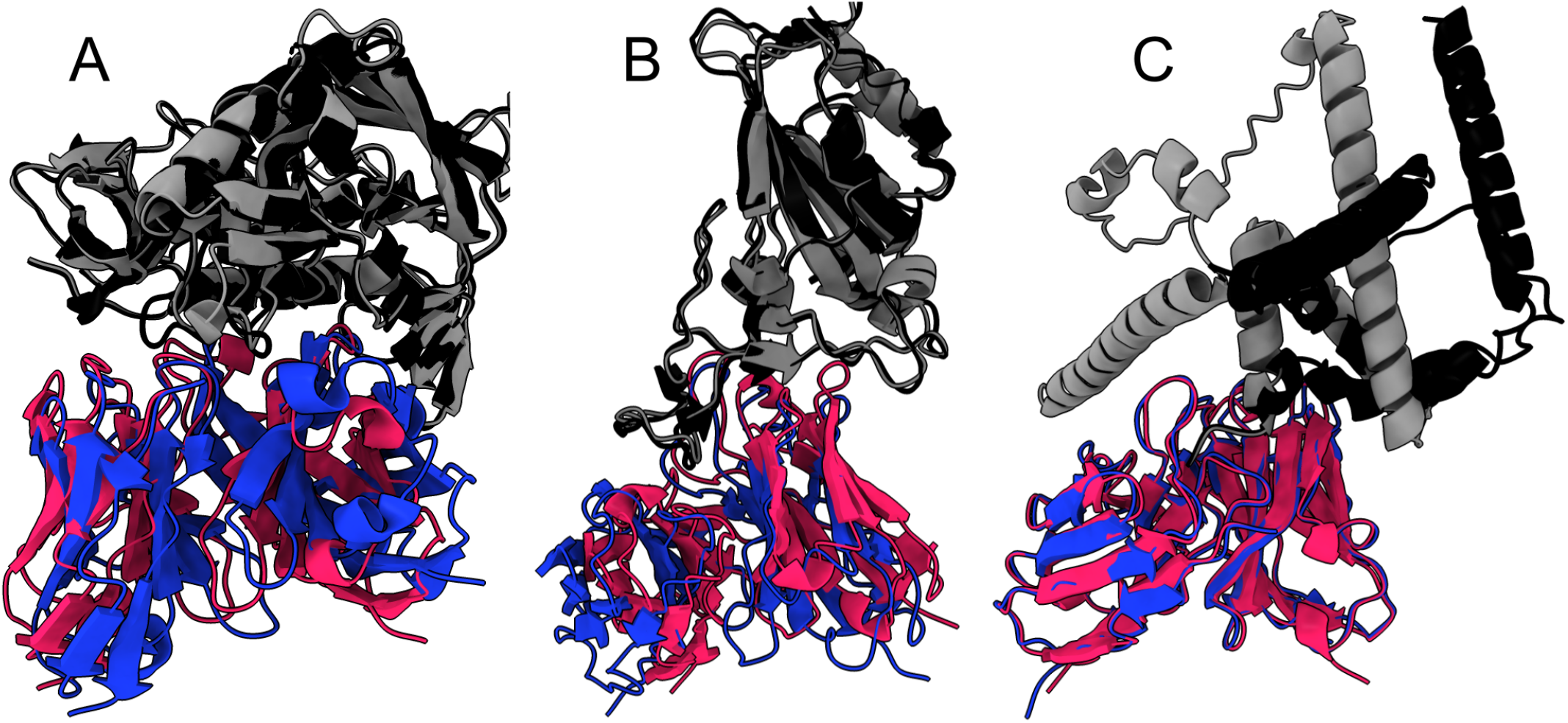
Examples of AF3 predicted true positives antibody-antigen complexes with low DockQ scores. Each image is zoomed in to focus on just the domain around the epitope. The ground truth antigen is colored black and ground truth antibody is colored blue. The AF3 antigen is colored gray and the AF3 predicted antibody in pink. The reference PDBs are (A) PDB: 5F9O, (B) PDB: 8KEO, and (C) PDB: 6CDE. Note that the antigen in (C) is modular – the different helices are connected by flexible linkers and this likely causes high RMSD in DockQ. Even though the immediate residues outside of the epitope do not superimpose cleanly, AlphaFold3 correctly detected the antibody should bind the tip of the helix.

We observed several false negatives that report small shift and displacement despite having poor DockQ scores (the points in Figure 7 (B) that lie near the y-axis). 1734 of the 1915 false negative complexes (roughly 90%) had a DockQ score less than 0.5; 651 of these 1915 false negatives (roughly 34%) had an epitope shift less than 10. We provide 3 examples of these in Figure 12. In these cases, AlphaFold3 detected the right epitope on the target, but failed to optimize the conformation of the interface, which increased the RMSD calculation driving a poor DockQ score. This is useful to know since additional tools (such as molecular dynamics) could be used to optimize the energy of these interfaces for complete evaluation, though this is beyond the scope of this project.

**Figure 12.**
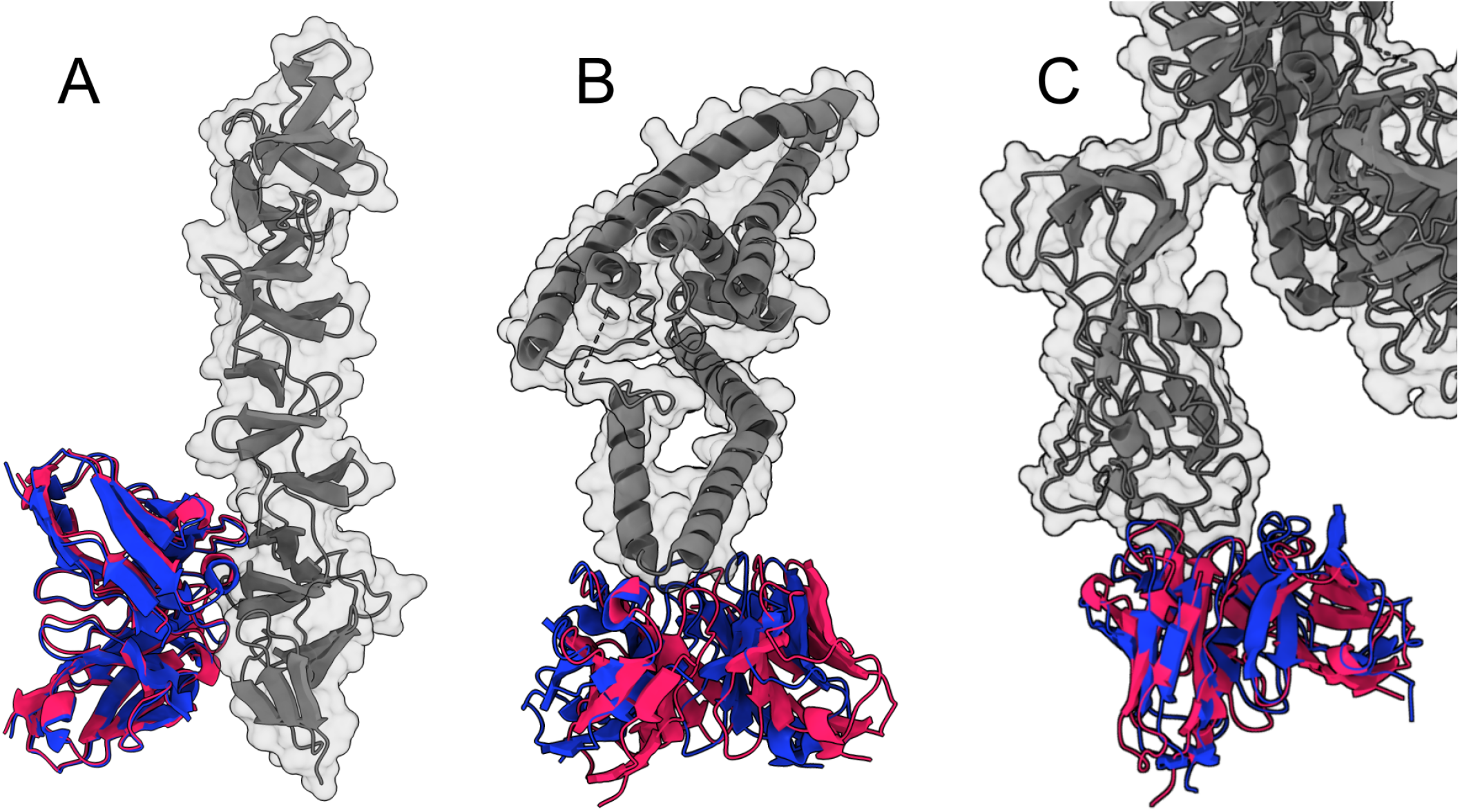
Examples of AF3 predicted false negative antibody-antigen complexes with small epitope shifts. Each image is zoomed in to focus on just the domain around the epitope. The true antigen from the PDB is colored gray and the true antibody is colored blue. The AF3 predicted antibody location is shown in pink. The reference PDBs are (A) PDB: 4NP4, (B) PDB: 7LJA, and (C) PDB: 7WTI.

We cannot evaluate the false positive epitopes predicted by AlphaFold3 with our current metrics. Methods such as DockQ and our algorithms rely on sequence alignment of the two structures – therefore we cannot compute the RMSD of two interfaces with unmatched targets. However, we can qualitatively assess the various epitopes on our Many-to-One batches A through F. For each of these batches, we collected all false positive complexes predicted by AlphaFold3, and superimposed these complexes on top of a reference model of its target protein using ChimeraX. This allowed us to measure the location of the false positive antibody relative to the target. Figure 13 shows the distribution of these false binders around the various target proteins. AlphaFold3 is capable of hallucinating epitopes all around the entire protein. Certain spots on the protein’s surface attract more antibodies, acting as “hotspots” for antibodies, but AlphaFold3 can scan the entire surface of a target protein and hallucinate binding signals anywhere on the globular domain of a protein.

**Figure 13.**
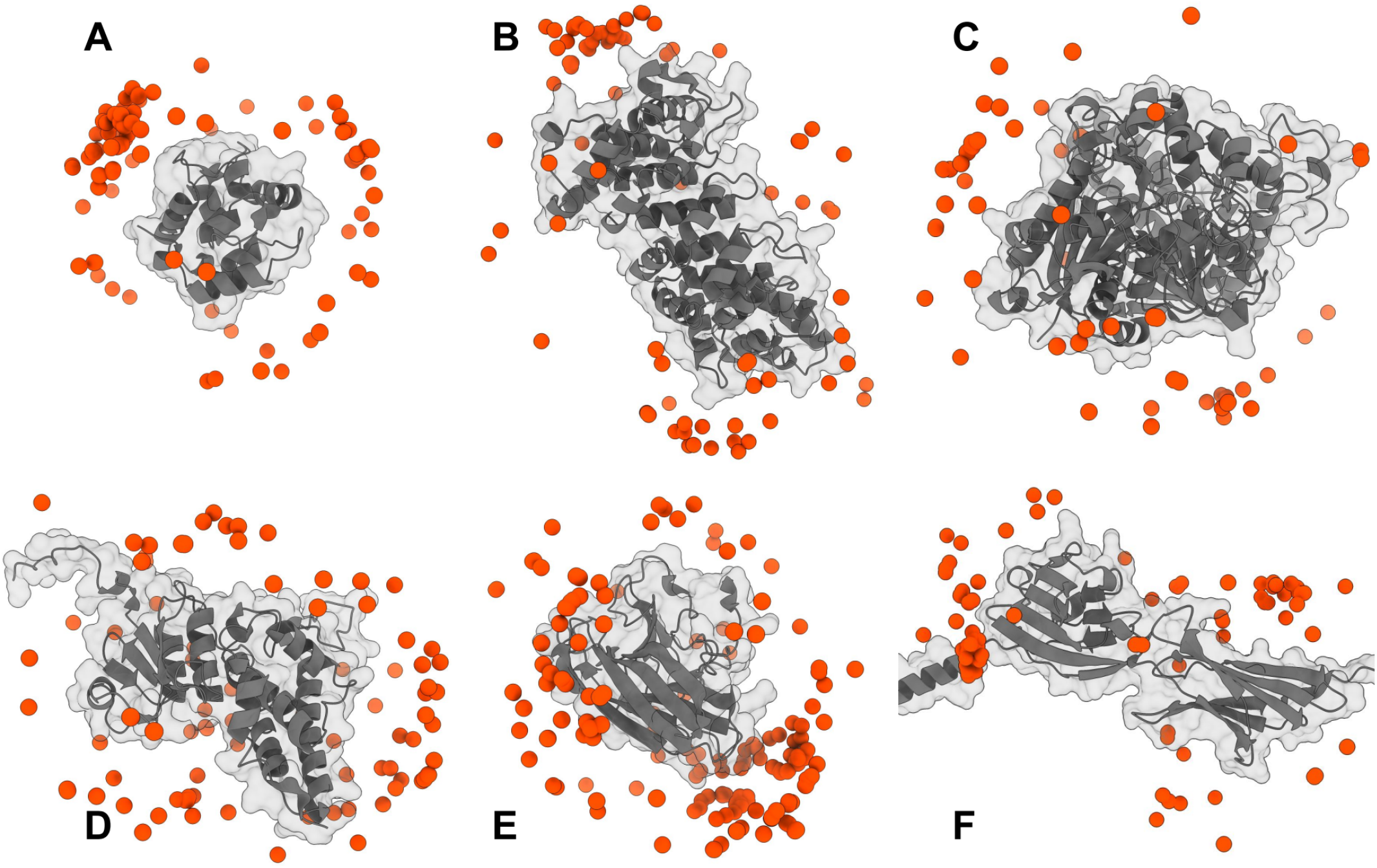
False positives around the Many-To-One proteins. Each target protein in the batches A through F is shown in gray. The center of each positively predicted antibody is shown as an orange sphere. AF3 can hallucinate false positives on diverse epitopes spanning nearly the entire surface of a well-folded protein. The disordered region is cropped off in (F).

## 5. Discussion

Our results reveal that several factors bias how accurately AlphaFold3 models antibody-target binding. The first is the source of the antibody and target structure – AF3 prefers X-ray crystallography derived PDBs over those derived from electron microscopy (EM). There are two possible explanations for this:

1. X-ray structures are overrepresented in the PDB, and thus dominate AF3’s training set compared to EM structures. Even though we noted in 4.3.1 that antibody leakage itself did not bias AlphaFold3, more X-ray PDBs did leaked through into our positive control dataset than EM PDBs. X-ray structures generally have better resolution than EM structures (16). In a sense this is a batch effect, and in our test we only provided AF3 with sequences to predict from, so this batch effect was likely learned by AF3 training.
2. The proteins captured in X-ray PDBs are different from those in EM PDBs. X-ray PDBs require the proteins of interest to be easy to crystallize for X-ray diffraction, while EM PDBs solve structures after freezing the proteins. This means EM proteins can be more flexible, disordered and dynamic while X-ray proteins are not. As a consequence, X-ray proteins are easier for AF3 to fold.

We believe a combination of both rationales, the batch effect and protein flexibility, contribute to the X-ray bias. One means of tackling this bias would be to test AF3 on a non-structural dataset: a dataset reporting antibody-antigen sequences only. Ideally such a dataset should also annotate the epitope too. We will investigate such datasets in future studies.

The X-ray bias points to the next challenge in structure and binding prediction: disorderness. The presence of disordered regions in a protein adds a “noise” in inference that can hinder epitope searching. Similar to how an unfolded protein cannot present a proper folded epitope, a disordered region could physically interfere with the antibody during prediction, or impact the nearby domains by changing their conformation. And in cases where an antibody is supposed to bind directly to the disordered region, AF3 mostly skip searching those regions and only focuses on well-folded domains. This is a consequence of disordered regions having low confidence in AF3 predictions.

An important observation in our study is that AF3 has an innate positive discovery rate. We generated false positives across all batches of our negative controls, showcasing how AlphaFold3 can hallucinate high quality interfaces that do not form in nature. We do not yet understand the mechanisms underlying this propensity – if it is driven by a particular residue motif or shape conformity, if it is a fictitious energy minima, or if there is some particular biochemical attribute causing this, is unknown.

The size and shape of the target protein are critical as well. Big proteins generate a large search space simply due to having more residues, and as such yield fewer positives. This can limit AF3’s utility in applications such as developing antibodies against novel targets. Proteins such as spike proteins on viruses can easily exceed over 1000 residues in length, and have oblong shapes with large surface areas. Screening antibodies that bind to such targets using AF3 would require more seeds and less stringent criteria, but this could yield more false positives while being computationally more expensive.

Lastly, AF3 predicted correct epitopes in several of our false negatives. Our epitope mapping highlighted how poor RMSD was the primary reason why these complexes had low confidence. Possible strategies to address this include the use of other tools such as molecular dynamics or docking to optimize these interfaces, increasing AF3’s recycling parameter to allow the model to refine the protein backbones and interface, and even finetuning AF3 for antibody-antigen prediction.

## 6. Conclusion

AlphaFold3 is a powerful tool that allows for the screening of antibodies against target proteins of interest with the mere sequences alone. In this study we conducted a comprehensive benchmark, testing AlphaFold3’s performance on a large antibody dataset and observed a maximum recall of 53% and a false positive rate of 3%. We noted how issues such as data leakage, intrinsic disorder of proteins, and large search spaces can impact AlphaFold3 predictions. Other factors such as the diversity of sequences encoded by the various CDRs of antibodies and the various possible epitope locations on a target do not seem to be a concern; AF3 does “model” any given antibody and target individually to predict their unique interface. It thoroughly scans the whole globular surface of the target protein for complementary epitopes. It is capable of accurately modeling antibodies and proteins it has never seen in training. Users should exercise caution when screening antibodies against proteins with more disordered regions and consider truncating the protein to reduce the complexity of their search.

We acknowledge that our analysis has several limitations. We only investigated the individual residues of the CDRs and their lengths, without the full context of the whole continuous sequence of each CDR. A more protein-language specific analysis, likely with protein language models could elucidate biases we may not have detected (25). Our epitope mapping scripts are also reliant on AF3 prediction quality - proteins with multiple domains can be tough to align through superimposition, and this can contribute to artifacts in our epitope shift and antibody displacement scores. A thorough analysis of the epitopes predicted by AF3 would also involve require analyzing the biochemistry properties of each epitope – the disorderness, hydrophobicity, charge distribution, etc. Since AlphaFold3 has biases towards PDB models, we believe benchmarking it over a sequence-only dataset could reveal further biases. Furthermore, in our study we did not test other protein folding tools on this same dataset as we wanted to start with AF3 due to its superlative accuracy (18; 15). We will expand our analysis to other folding models such as Boltz-2, RoseTTAFold2, and Chai-1 in the future (18; 23; 3). Finally, our immediate priority for future inquiry is understanding what differentiates the hallucinated interfaces of false positives from true antibody-target interfaces, as this would be immensely valuable in digital antibody screening.

## 9. Data availability

The data used to conduct this study can be freely accessed from SAbDab, UniProt, and AlphaFold Protein Structure Database. The data generated over the course of our analysis is currently being withheld due to ongoing research. However, our methods delineate how readers can construct their own datasets.

## 10. Author contributions statement

AS, ZW and WJZ conceived the project. AS, NSM, MR, RL, WC, ZW and WJZ contributed to experiment design. AS and NSM conducted the experiments. AS analyzed the results. AS and WJZ wrote the manuscript.

## A. Detailed Methods

### A.1. Data Mining

In May 2025, we downloaded 3096 PDB files and their corresponding FASTA files that contained antibodies binding single-chain protein targets from SAbDab. We extracted 3401 non-redundant amino acid sequences of the antibody heavy chains, light chains and the antigen from these files. Some of the PDB files included multiple antibodies bound to the same target. We restricted the sequences of the antibody chains to just the variable fragment, i.e. position H113 on the heavy chain and L107 on the light chain, using the chothia numbering provided by SAbDab for each antibody chain.

For every target protein longer than 500 residues we attempted to fragment the target to isolate the subunit binding the antibody. By using the PDB file, we sliced the target if a single subunit with a continuous sequence was contained within 20 angstrom of any antibody atom. The rationale for this was to focus on the target residues involved in binding. Only 133 targets could be successfully fragmented, while 672 other targets had to be saved with their full sequences.

Alongside the sequence of the three chains, we also tracked other information such as the position and sequence of the CDRs, the original structure in the PDB, the date of the PDB’s publication, and most importantly the type of experiment used for deriving the PDB. Our dataset contained 1342 sequence triplets derived from Electron Microscopy, and the remaining 2059 were captured from X-ray Crystallography. These 3401 datapoints form our positive control in this study. To observe if Alphafold3 prefers one type of experimental method over another, we labeled all complexes as either M if they were derived from PDBs solved using electron microscopy, or X from X-ray crystallography.

One out of the 3096 PDBs that we downloaded (PDB: 4CC8) had improper residues. All target residues in this PDB were masked as X. We manually extracted the sequence of the claimed target (“HIV-1 envelope glycoprotein”) from UniProtKB.

#### A.1.1. Complementarity Determining Region Definitions

SAbDab provides a chothia-indexed file with sequences of all antibody chains it contains. Using these indexes, we identified the CDRs of all 3401 antibodies with our own definitions (as listed in Table 3) to capture all potentially relevant residues. Our CDR definitions are based off of previously published definitions (such as Chothia, Kabat and IMGT (7; 1; 8; 9)), but span the aggregate length of CDRs covered by these definitions. That is, our definitions are a union of these definitions. We measured the length of all CDRs in our dataset, and also compiled each individual amino acid constituting them (Figure 6).

#### A.1.2. One-to-One Negative Targets

We downloaded the sequences of all human proteins between 50 to 500 residues in length from UniProtKB. For each of the 3401 positive control antibodies, we paired a randomly sampled human protein after verifying the protein had low sequence similarity to the true antigen. Note that these targets were extracted from UniProt, a separate database than SAbDab, and this introduced a potential confounding factor in our antibody-target screening using AlphaFold3 – before modeling the binding complex of the antibody and negative target, AlphaFold3 had to first “solve” the structure of the target. Since these proteins were sampled by their sequence, it is possible that several of these targets could have weak structure predictions due to higher disorderness. This is the rationale behind why we generated more negative control data points than just the One-to-One negative targets.

#### A.1.3. Many-to-One Negative Target

Here we list details about the 6 proteins we selected to form our negative control batches A through F. Their high quality structures as per AlphaFold Protein Structure Database are noted in Table 4.

1. *Rattus norvegicus* PVALB: a 110 residue long protein, with no disordered regions and several experimental structures in the PDB. None of these PDBs contained antibodies binding it.
2. *Drosophila melanogaster* CG6073: a 369 residues long protein, with no disordered regions and no experimental structures in the PDB.
3. *Escherichia coli* araB: a 566 residue long protein, with a small (10 residues long) N-terminus peptide region and no experimental structures in the PDB.
4. *Bos taurus* CLIC4: a 253 residue long protein, with a small (10 residues long) C-terminus peptide region and no experimental structures in the PDB.
5. *Arabidopsis thaliana* DGR2: a 186 residue long protein, with no disordered regions and no experimental structures in the PDB.
6. *Homo sapiens* CD274: a 290 residue long protein with a globular shape flanked by two alpha-helices (one on each teminus). This protein, also known as PD-L1, is the target of several therapeutic antibodies and has many PDBs, several of these containing antibodies binding it. A total of 9 antibodies in our positive control were derived from PDBs with PD-L1, so we removed these 9 antibodies from this batch of negative control. The rationale for picking this target was to investigate a well studied cancer-related protein and observe any biases AlphaFold3 had towards it.

We ensured the Many-to-One targets that we manually picked had properly folded shapes with few disordered regions. Table 4 lists their high quality structures as per AlphaFold Protein Structure Database. For all predictions involving a Many-to-One negative target, we used the PDB file from AlphaFold Protein Structure Database as a template to ensure we predict binding on the same backbone and did not “burden” AF3 with extraneous folding.

### A.2. AlphaFold3 Screening

We installed AlphaFold3 locally on our servers and predicted the complexes of all data points. We ran the default AlphaFold3 data pipeline (this includes the sequence alignment and template searching) on a CPU server and then inferred the structures for 10 seeds on a GPU-intensive server. We also predicted the structures of the lone target separately. We collected all the confidence scores generated by AlphaFold3 in the main summary file such as predicted aligned error (PAE), interface predicted template modeling score (IPTM) and ranking score.

In our analysis, we calculated the PAE score between the antibody and target as the minimum PAE out of the 4 possible interface combinations between the antibody and the target: heavy chain to target, light chain to target, target to heavy chain, and target to light chain. Note that despite PAE being a per-residue pair error measurement, it is not perfectly commutative as a confidence metric. That is, the PAE score of some residue *i* to another residue *j* is not the exact same value as that from *j* to *i*. The IPTM score we used was the mean of the IPTM scores from the target to the heavy chain, and from the target to the light chain.

For our Many-To-One negative controls, we used the AlphaFold Protein Structure Database models as the only templates for AlphaFold3 inference. We did this to ensure that exactly the same input conditions were used by AlphaFold3 to fold those target proteins across different antibodies.

### A.3. Epitope Mapping

We assumed that since all antibodies in our dataset had been successfully imaged in the PDB, they must have had mature affinity to their true antigen.

#### A.3.1. DockQ

In our study, we calculated the DockQ score of a predicted complex as the mean of the DockQ scores of the heavy chain to the target and the light chain to the target. Additionally, we modified the DockQ source code in our local environment to force it to compute on all interfaces; the original DockQ threshold would ignore several of the AlphaFold3 predicted complexes due to not detecting interfaces. As expected, all of these forced interfaces produced low DockQ scores. As DockQ correlates with the RMSD of the antibody-target interface, we reasoned that a low DockQ score for a complex would not guarantee that the correct epitope was not detected. In instances where the predicted complex had a Low DockQ score, AlphaFold3 could have predicted the correct epitope but still produce poor RMSD alignment. This is the rationale behind why we used our own epitope metrics alongside DockQ. Using all three metrics, we could more accurately assess what exactly occurred in any AlphaFold3 prediction.

#### A.3.2. Epitope Shift

Epitope shift tracks the location of the true and predicted epitopes for an antibody-target complex, without requiring any structure superimposition. For this script, we define the epitope as any residue on the target that has an atom within a 4 angstrom distance of any atom in the antibody. We first identify the epitope *G* in the ground truth PDB, and then the epitope *P* in the predicted model. From the predicted model we measure the coordinates of the *α*-Carbon of every residue of the target. Using these coordinates, we then compile the backbone locations of both epitopes *G* and *P*, and then calculate the absolute difference in the mean of both sets. We term this difference as “Epitope Shift”, and it measures how much the epitope “shifted” from the ground truth to the predicted model. A small epitope shift indicates that the sets *G* and *P* consist of the same residues and that the predicted model features the true epitope.

#### A3.3. Antibody Displacement

Antibody Displacement measures the change in the predicted antibody’s position relative to the ground truth position. For this algorithm, we used ChimeraX to superimpose the predicted antibody-antigen model to the ground truth PDB. Note we only superimpose using the antigen protein’s residues, i.e. the RMSD optimization ignores the antibody completely when minimizing. After the superimposition, we measure the center of mass of the first 130 residues of the heavy chain of the antibody for both the ground truth and predicted model. We calculate the absolute difference of these two centers as the “antibody displacement”. A low displacement from the ground truth to the prediction means that the relative location of the antibody with respect to the antigen is unchanged.

### B. More Statistical Results

#### B.1. AlphaFold3 confidence correlation

We used the Mann-Whitney U test from SciPy Statistics to compute the significance of each confidence score generated by AlphaFold3 for an antibody-target complex. We split the set of 27199 tested complexes into the positive and negative controls, and computed the p-value of the difference of each confidence score across these sets. In order of decreasing significance, these are:

1. Chain-pair minimum PAE Heavy to Target (p-value = 1.24 *×* 10^−307^)
2. Chain-pair minimum PAE Light to Target (p-value = 1.99*×* 10^−300^)
3. IPTM (p-value = 1.19 *×* 10^−287^)
4. Ranking Score (p-value = 5.60 *×* 10^−261^)
5. Chain PTM Light (p-value = 2.83 *×* 10^−41^)
6. Chain-pair IPTM Heavy to Heavy (p-value = 2.83*×* 10^−41^)
7. Fraction disordered (p-value = 3.57 *×* 10^−17^)
8. Chain PTM Target (p-value = 9.43 *×* 10^−10^)
9. Chain-pair IPTM Target to Target (p-value = 9.43*×*10^−10^)
10. Ranking Score (p-value = 5.60 *×* 10^−261^)

The remaining metrics had p-values greater than 0.05, and are not listed here due to lack of significance.

### B.2. Target Measurements

Table 5 lists the number of complexes from the M, X, and ℕ in each bin. The trend of decreasing positive rate with larger size was remained when we removed the negative controls.

## C. More Epitope Results

Figure 11 shows examples of true positive predictions that had DockQ scores less than 0.5, but also had epitope shifts less than 10 angstrom. These are complexes were AlphaFold3 predicted the correct epitope and had high confidence, but did not predict the optimal conformation of the interface leading to high RMSD.

Figure 12 shows examples of false negative predictions that had DockQ scores less than 0.5, but also had epitope shifts less than 10 angstrom. These are complexes in which AlphaFold3 predicted the correct epitope but with low confidence and high RMSD.

Figure 13 shows all false positives predicted on the Many-to-One batches.

